# Atlas of transcriptionally active transposable elements in human adult tissues

**DOI:** 10.1101/714212

**Authors:** Gireesh K. Bogu, Ferran Reverter, Marc A. Marti-Renom, Michael P. Snyder, Roderic Guigó

## Abstract

Approximately half of the human genome consists of mobile repetitive DNA sequences known as transposable elements (TEs). They are usually silenced by epigenetic mechanisms, but a few are known to escape silencing at embryonic stages, affecting early human development by regulating nearby protein-coding genes. To investigate transcriptional activity in human adult tissues we systematically investigate the expression landscape of about 4.2 million non-coding TEs in 8,051 RNA-Seq datasets from up to 49 adult tissues and 540 individuals. We show that approximately 79,558 individual TEs (2%). belonging to 856 subfamilies escape epigenetic silencing in adult tissues and become transcriptionally active, often in a very tissue-specific manner. Supporting a role for TEs in the regulation of expression of nearby genes, we found the expression of TEs often correlated with the expression of nearby genes, and significantly stronger when the TEs include eQTLs for the genes. We identified thousands of tissue-elevated, sex-associated TEs in the breast, ethnicity-associated in the skin and age-associated in the tibial artery, where we found a potential implication of two TE subfamilies in atherosclerosis. Our results suggest a functional role of TEs in the regulation of gene expression, support their implication in human phenotypes, and also serve as a comprehensive resource of transcriptionally active TEs in human adult tissues.

## Introduction

Transposable elements (TEs) are widespread self-replicating non-coding repetitive DNA sequences occupying more than half of the human genome sequence. They have been shown to be globally repressed by various epigenetics mechanisms especially DNA methylation (Chuong et al. 2017). However, recent transcriptomic studies have shown that several subfamilies of TEs are expressed in human tissues, mostly at embryonic stages (Göke et al. 2015; Wang et al. 2014; Grow et al. 2015). It is unclear whether the same trend characterizes adult human tissues.

TEs can be categorized into two broad types – retrotransposons and DNA transposons. Retrotransposons spread by a copy-and-paste mechanism with the help of their RNAs, whereas DNA transposons spread directly without an RNA intermediate (Elbarbary et al. 2016). Transposition of retrotransposons is highly active in early embryonic tissues and also in the adult brain (Erwin et al. 2014). TEs harbor transcription factors binding sites and can act as transcriptional enhancers, promoters, and silencers and thereby regulate neighboring gene expression (Chuong et al. 2017). The functional activity of TEs is not only exerted at the DNA level but also at the RNA level. A well-known example is the X-chromosome inactivation where silencing of the X-chromosome is mediated in part by RNA intermediates from L1 sequences (Elbarbary et al. 2016). A recent study has shown that RNA from LINE1 sequences mediate binding of nucleolin and Kap1 to ribosomal DNA promoting ESC self-renewal (Percharde et al. 2018).

Previous transcriptome studies identified several transcriptionally active TEs in humans. An initial genome-wide retrotransposon transcriptome study using CAP Analysis Gene Expression Sequencing (CAGE-seq) data from 12 tissues identified 23,000 candidate regulatory regions derived from retrotransposons (Faulkner et al. 2009). A follow-up CAGE-seq study using both nuclear and cytoplasmic transcriptome data from human embryonic stem cells, induced pluripotent stem cells, fibroblasts, and lymphocytes showed that subfamilies of endogenous retroviruses (ERV1 family of LTR class) including LTR7, HERVH-int and LTR9 were highly expressed TEs in human stem cells (Fort et al. 2014). RNA-seq data in human embryonic and pluripotent stem cells revealed similar trends, where the HERVH subfamily is highly expressed and marked with the active promoter and enhancer chromatin marks (Wang et al. 2016). Furthermore, a recent single-cell RNA-seq study also revealed the higher expression of ERVs at various stages of preimplantation embryogenesis in human and showed their ability to distinguish all cell-specific stages using TE transcription alone (Göke et al. 2015). This study also demonstrated that TEs play a significant role in fine-tuning cellular functions in early human development (Grow et al. 2015; Göke et al. 2015) by controlling neighboring gene expression.

Different factors such as stress and heat shock have been shown to activate transcription of many TEs in humans (Häsler and Strub 2006). Transcriptional activation of TEs in the context of senescence and aging in human fibroblasts, mouse tissues, and fly brain also have been reported (De Cecco et al. 2013b, 2013a; Li et al. 2013). Notably, L1 subfamilies were shown to be upregulated in 36 month-old mouse liver compared to 5 month-old (De Cecco et al. 2013b). Moreover, mRNA expression has been shown to decrease with age (De Cecco et al. 2013b). In the fly brain, TE transcriptional activation during normal aging has shown to be associated with neuronal decline suggesting the functional significance of TEs in the aging process (Li et al. 2013).

Analysis of the pilot GTEx (Genotype-Tissue Expression) data (Melé et al. 2015) (1,486 RNA-seq samples from 175 individuals) provided initial support for the transcriptional activation of TEs in human adult tissues. Here, we analyzed a much larger GTEx dataset to systematically investigate the general transcriptional patterns of TEs and how they correlate with human phenotypes. We used very stringent criteria to identify TEs likely to be *bona fide* independent transcriptional events and not a part of larger transcriptional units. We uncovered extensive, and largely tissue-specific, TE transcription, and found evidence role for TEs in regulating gene expression of nearby protein-coding genes. We also found associations between tissue-specific TE expression and a number of human phenotypes. Thus, we found female breast expressing a much larger number of TEs than male breast, as well as the skin of African Americans, compared to the skin of individuals of European Ancestry. We found a substantial increase of TE expression with age in a number of neural and non-neural tissues, but most notably in the tibial artery, and found a potential implication of a few TE subfamilies in atherosclerosis. In summary, we create the most exhaustive atlas to date of TE expression in human adult tissues.

## Results

### Overall approach

To systematically examine the transcriptionally active TEs, we analyzed RNA-seq data derived from 8,051 samples from 49 tissue-sites of 540 post-mortem donors, corresponding to the version v6 of GTEx (GTEx Consortium et al. 2017) (Fig. 1).

**Figure 1.**
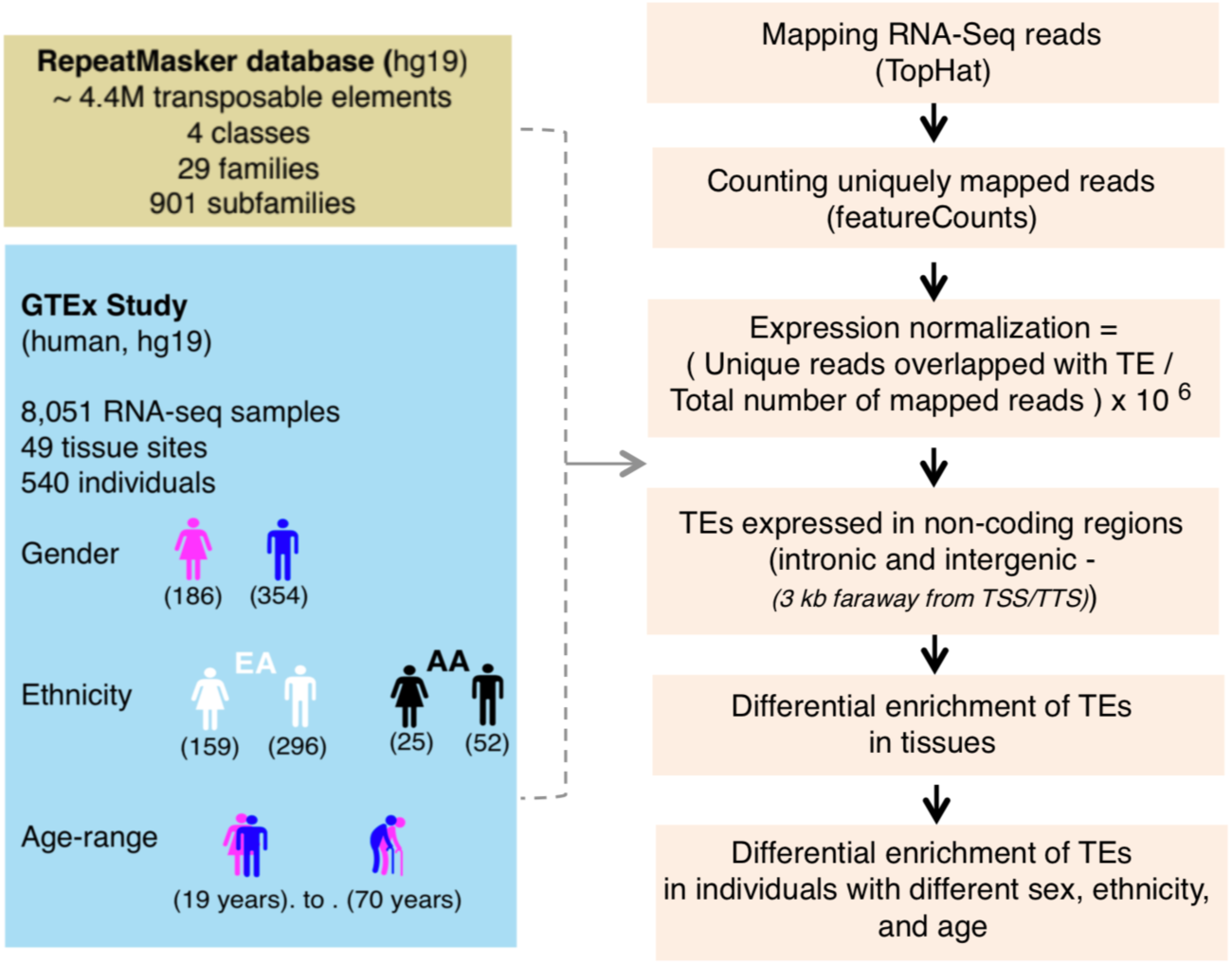
Study Overview. Transposable elements (TEs) expression was quantified by using the individual TE genomic copies (around 4.4 million individual copies were grouped into 4 classes or 29 families or 901 subfamilies) from RepeatMasker database (hg19) and 8,051 RNA-seq datasets from GTEx (Genotype-Tissue specific Expression) study. 4.2 million copies occurred in non-coding regions of the genome. Expression of TEs was quantified by using only uniquely mapped reads. TEs overlapping exons and regions near (within 3 kb) transcription start site (TSS) or transcription termination site (TTS) of protein-coding genes were removed. Differential enrichment of TE expression was analyzed in tissues and individuals with different sex, ethnicity, and age.

Using this data, we examined the expression of 4.2 million individual TE sequences (grouped into 901 subfamilies (Smit et al. 2010)) occurring in non-coding regions (introns and intergenic regions). We applied a number of filters to the mapped reads in order to obtain robust estimates of expression of TEs (Fig. 1, **Methods**). In particular, to rule out that expression of TEs is a by-product of the pervasive transcription of nearby genes, we took a conservative approach, generating transcriptional peaks from RNA-seq reads and considered a TE expressed only when overlapping one such peak (**Supplemental Fig. 1A, B, C).** Thus we did not consider a TE expressed if embedded in a larger transcriptionally active region.

### TE expression is associated with tissue type

In total, we found 79,558 TE elements in non-coding regions (about 2% of all non-coding TEs) expressed in at least one tissue sample (**Supplemental Table 1**), and these belonged to 856 subfamilies (95% of all subfamilies) (**Methods**). Unsupervised clustering of 8,551 samples using expression data with 79,558 TEs reflected tissue groups. Despite the exclusion of TEs that were either in exonic regions or proximal to protein-coding genes, tissues-types originating from the same tissue often aggregated by the tissue-specific expression, supporting, a possible biological role for TE expression in defining tissue type (Fig. 2A, **Methods**).

**Figure 2.**
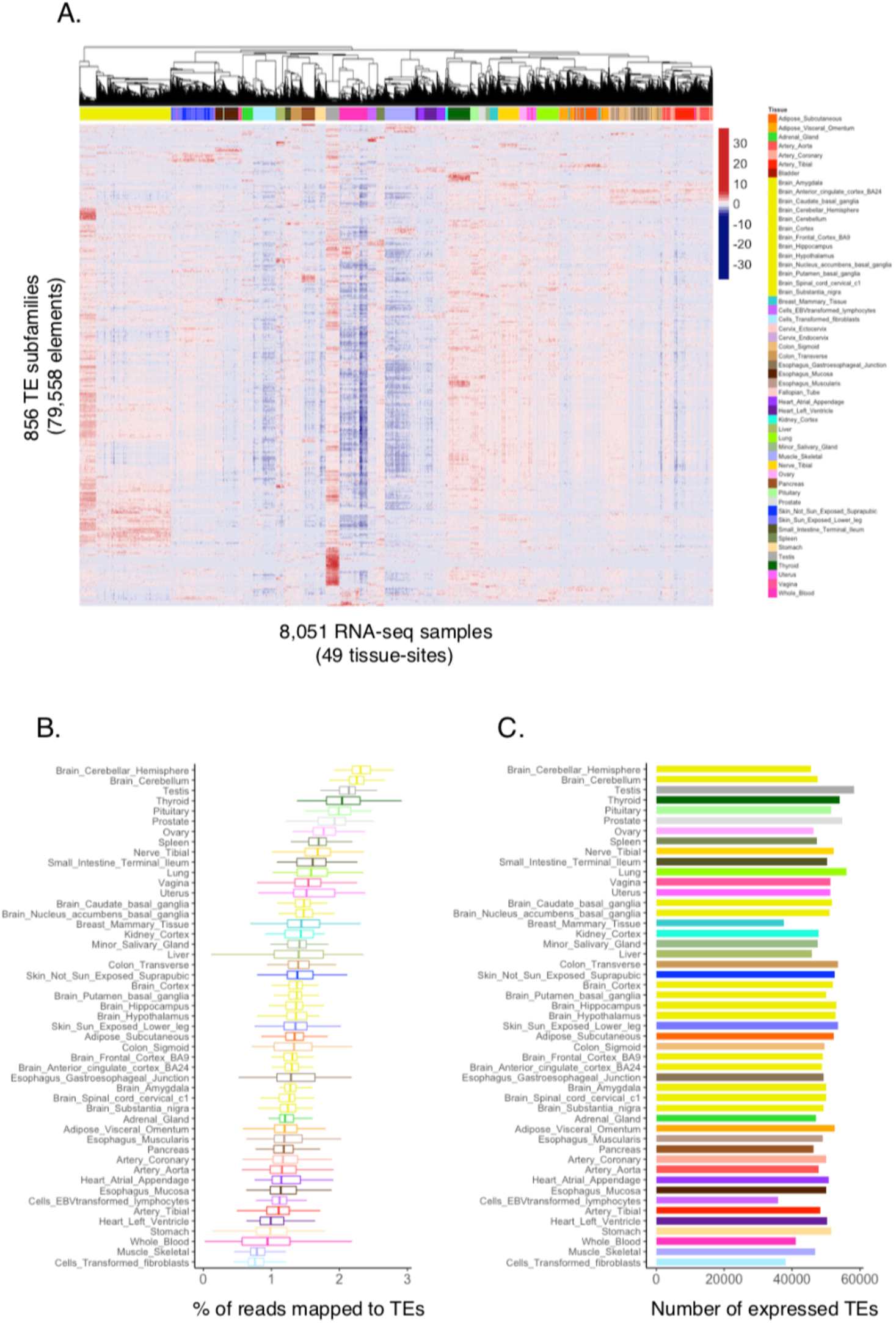
Expression of transposable elements. (**A**) Unsupervised clustering of TE subfamilies on the basis of their expression recapitulates tissue types. 856 tissue-specific subfamilies were shown in rows and 8,051 RNA-seq samples in columns. Clustering bar on the top of the heatmap with different colors labels the different tissues (**B**) Boxplots show the distribution of the percentage of reads that mapped to TEs across different tissues (Percentage was calculated by dividing the number of uniquely mapped reads overlapping TEs with the total number of mapped reads in each sample and further multiplied by a million). (**C**) Number of TEs expressed in multiple adult human tissues (To estimate the number of individual genomic TE copies expressed in each tissue, we counted the ones that have greater than or equal to 0.1 RPKM expression in at least one sample in each tissue separately).

While we did not find large differences in the number of expressed TEs across tissues, the overall TE expression, measured as the proportion of reads mapping to TEs, varied several-fold, with cerebellum and testis showing the highest percentage of TE transcription, and transformed fibroblasts and skeletal muscle the lowest (Fig. 2B, C).

### TEs may contribute to regulating expression of nearby protein-coding genes

We examined whether TEs could play any role in regulating the expression of nearby protein-coding genes. To address this, we overlapped eQTLs from a recent GTEx study (GTEx Consortium et al. 2017) pooled from all tissues with our non-coding TEs. We found a significant enrichment (Fisher’s exact test, P-value < 0.05) of expressed TEs (25%) overlapping eQTLs compared to non-expressed (16.6%) (Fig. 3A, **Supplemental Table 2**) indicating a potential regulatory role of TEs.

**Figure 3.**
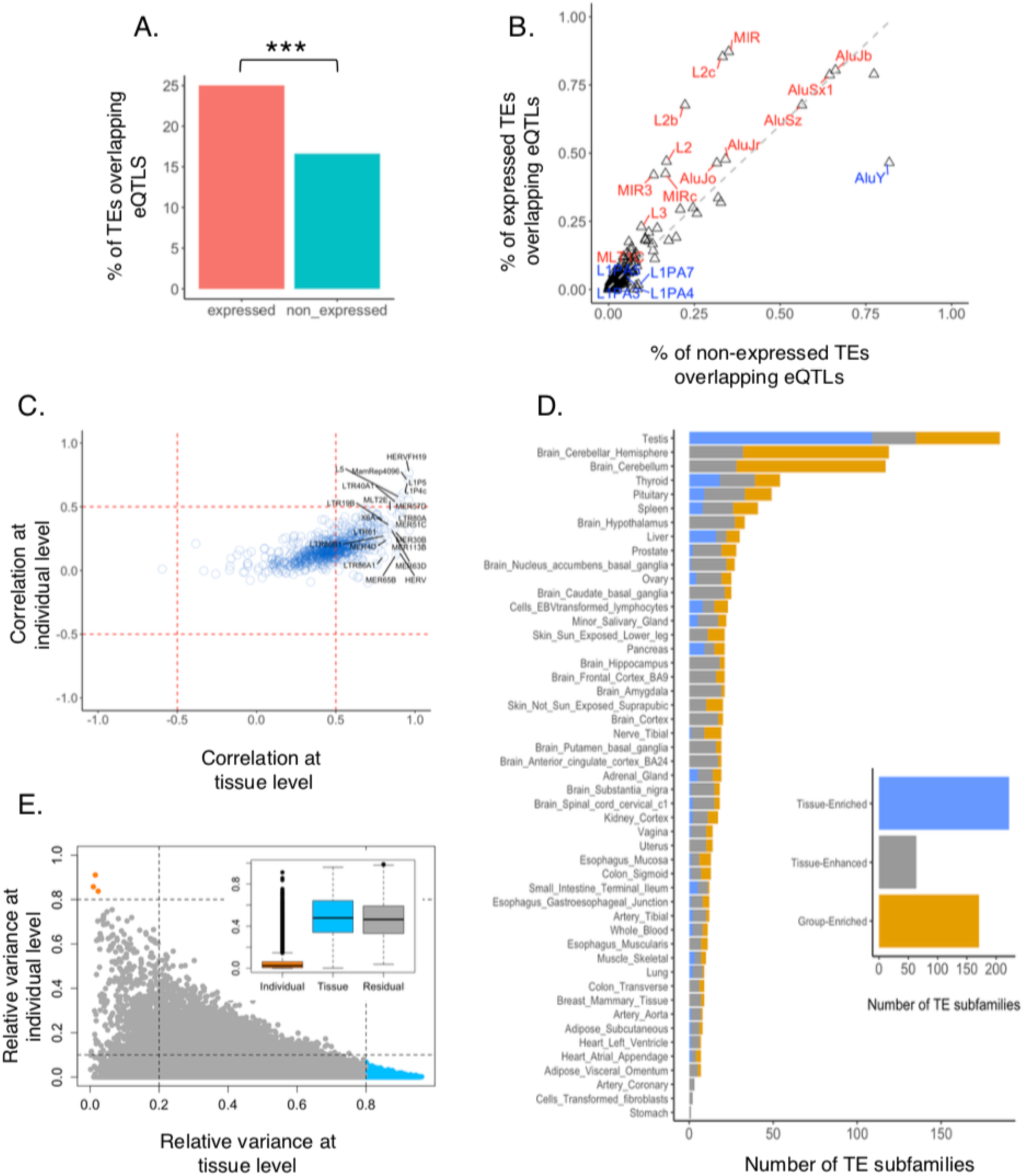
Potential regulatory role of transposable elements and variance decomposition of TE expression across tissues and individuals. (**A**) Percentage of TEs overlapping eQTLs across 49 tissues. 25% of expressed TEs overlap eQTLs compared 17% of non-expressed TEs (Fisher-exact test, *** = p-value < 0.05). (**B**) Percentage of TE subfamilies overlapping eQTLS across 49 tissues. The x-axis represents non-expressed subfamilies and Y-axis represent expressed. Enriched subfamilies were highlighted in red and depleted in blue. (**C**) Correlation between TEs and their nearest protein-coding genes in tissues and individuals. Each circle represents the median correlation within TE subfamilies. Correlations were mostly at the tissue level and in a positive direction. Top right and middle right quadrants represent a few examples of the top TE subfamilies that were positively correlated at the tissue level. (**D**) Differential enrichment of TEs across tissues. Right: Stacked bar plot shows 222 tissue enriched subfamilies in blue (at least 1.5 fold higher expression levels in a particular tissue as compared to all other tissues), 171 group-enriched subfamilies in gold (at least 1.5 fold higher expression levels in a group of 2-7 tissues) and 64 tissue enhanced subfamilies in grey (at least 1.5 fold higher expression levels in a particular tissue as compared to average levels in all tissues) colors. Left: Tissue-enriched, group-enriched and tissue-enhanced subfamilies separated by tissues. (**E**) The contribution of tissue and individual to the variance of TE expression. Bottom right: TEs with high tissue variation and low individual variation. Top Left: TEs with high individual variation and low tissue variation. Inset: Boxplot showing the contribution of variance across individuals and across tissues to the total variance in TE expression

We explored whether the enrichment varies across different TE subfamilies. Among the subfamilies most highly enriched for eQTLs, we found L2, L2b, L2c of LINE class and MIR of SINEs (Fig. 3B). Our results are in agreement with a previous study of lymphoblastoid cells (Lappalainen et al. 2013), and suggests that TEs can harbor regulatory elements of nearby genes in human, the regulatory effect of which may be mediated by TE expression.

To further investigate the potential role of TEs regulating the expression of nearby protein-coding genes, we computed the correlation between the expression of TEs and that of the closest gene. To account for the fact that this correlation has two components, which are not necessarily in the same direction: correlation across tissues, and across individuals, we applied a multivariate multilevel model (MVML) to decompose the correlation between TE and gene expression pairs at the tissue and individual level (**Methods**). Using this model, we found the correlation dominated by correlation across tissues rather than across individuals, and more often positive than negative, in particular across individuals (Fig. 3C, **Supplemental Fig. 2A)**. Supporting a role for TEs in regulating gene expression, we found a correlation between a TE, and its closest protein-coding gene to be overall more positive stronger when the TE included an eQTL for the gene (0.50 vs 0.57, Wilcoxon test, P-value < 0.001, **Supplemental Fig. 2B, Methods**), and stronger both for positive (0.61 vs 0.58) and negative correlation (−0.20 vs −0.17).

### Extensive tissue-elevated TE expression

Based on the pattern of TE expression across all tissues, all 856 expressed TE subfamilies were classified into two major groups, 398 housekeeping (expressed in a similar manner across all tissues) and 457 tissue-elevated (expressed in a differential manner) (**Supplemental Table 4, Methods**).

Tissue-elevated subfamilies were further classified into three subgroups, 222 tissue-enriched (at least 1.5 fold higher expression levels in a particular tissue as compared to all other tissues), 171 group-enriched (at least 1.5 fold higher expression levels in a group of 2-7 tissues) and 64 tissue-enhanced (at least 1.5 fold higher expression levels in a particular tissue as compared to average levels in all tissues) (Fig. 3D, **Supplemental Table 4**). Testis showed the highest number of tissue-enriched and the cerebellar sections of the brain showed the highest number of group-enriched TE subfamilies (Fig. 3D, **Supplemental Figure. 3. A,B**). For example, LTR75B was a group-enriched subfamily as it was highly enriched in several brain tissue groups (**Supplemental Figure. 3C**), and, tissue-enriched subfamilies like LTR88c and LTR28 were enriched only in the pancreas and pituitary, respectively (**Supplemental Figure. 3C**). At the elemental level, we found 32,857 tissue-enriched and 18,895 group-enriched TEs (**Supplemental Table 4**), revealing a widespread amount of tissue-elevated TE expression in humans.

Tissue-enriched TEs were associated with xenobiotic, hormone and fatty acid metabolic processes (**Supplemental Table 4**). The link between xenobiotic processes and TEs transcription was previously observed in an epigenetic-drugs induced TE-transcriptome study (Brocks et al. 2017). Group-enriched TEs were associated with several biological functions including chromatid organization and segregation, amino acid transport, and amyloid precursor protein and fatty acid metabolic processes (**Supplemental Table 4**). Taken together, TE expression enrichment in various tissues and their association with a different biological process are consistent with a possible biological role of TEs in human adult tissues.

### Sex-associated TE transcription is most prevalent in the breast

Using linear mixed models (**Methods**), we found that variation in TE expression is far greater among tissues (48% of the total variance in TE expression) than individuals (2% of the total variance, Fig. 3E, **Supplemental Table 5**). The pattern of variation of TE expression across tissues and individuals is nearly identical to that reported for protein-coding genes (47% and 4%, respectively (Melé et al. 2015)). Gene ontology analysis of the protein-coding genes nearby TE the expression of which varies a lot across tissues, but little across individuals, indicates that these genes are involved in the regulation of neural activity (**Supplemental Figure. 3. C)**, consistent with the strong transcriptional divide separating neural from non-neural tissues. We investigated a number of phenotypes that could explain the variation of TE expression across individuals. Specifically, we investigated differential TE expression with respect to sex, ethnicity, and age.

We use the linear models above on tissues from 186 females and 354 males (**Supplemental Figure. 4**), and identified first 647 subfamilies (103 increased in male, 584 increased in female) that showed significant differential expression in at least one tissue (adjusted P-value < 0.05, **Methods**) between male and females (Fig. 4A, **Supplemental Table 6**). At element level, we identified 23,462 differentially expressed TEs (4,758 increased in male, 18,704 increased in female, **Supplemental Table 6**). The size effect, however, was small (in all cases, log2 fold-change < 1) (Fig. 4A).

**Figure 4.**
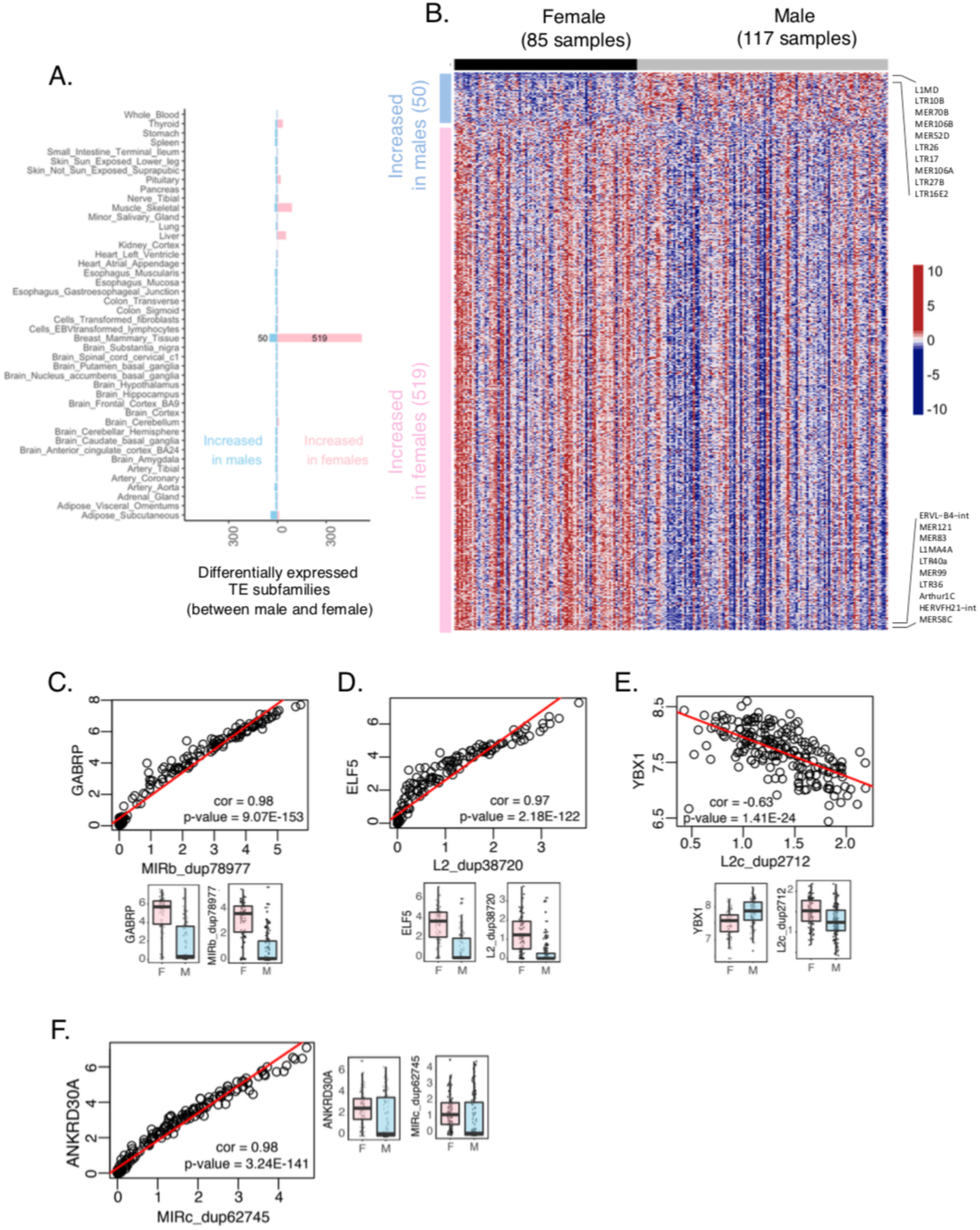
Differentially expressed TE subfamilies between males and females. (**A**) Double-sided stacked bar plot represents the number of differentially expressed TE subfamilies between males and females across different tissues. TEs the expression of which increased in males are shown on the left side and those with increased expression in females on the right. The highest number of differentially expressed TE subfamilies were found in breast mammary tissue (569, log2 fold-change > 0 or < 0, adjusted P-value < 0.05) and most of them were up-regulated in females. (**B**) Heatmap represents the expression of differentially expressed TE subfamilies in breast mammary tissue (red is higher and blue is lower). 569 subfamilies were shown in rows and 241 breast mammary samples in columns. Bar with the pink color on the left side of the heatmap represents a group of subfamilies that were increased in females and the one with the light blue color represents that were increased in males. Black and grey bars on the top of the heatmap identify the female and male samples. (**C-F**) Examples of sex-associated TEs that were correlated with stress fiber associated protein-coding genes in the breast mammary tissue. Scatterplots showing the TE expression (log2) on X-axis and gene expression (log2) on Y-axis, and the red line is the regression line. Each scatterplot is further separated into two (gene and TE) individual boxplots, where log-transformed expression was plotted in females and males. Females were shown in pink and males were shown in light blue colors. (**C**) For example, GABRP was positively correlated with a MIRc_dup78977 copy in breast mammary tissue where both GABRP expression and MIRc_dup78977 expression higher in females. (**D-F**) Similar plots were drawn using other top correlated TE and stress fiber associated gene pairs.

Breast mammary tissue exhibited the largest number of differentially expressed TEs (569 subfamilies, 16,732 elements). The vast majority were increased in females (519 subfamilies), mirroring sex-biased protein-coding gene expression (Melé et al. 2015) (Fig 4A, B). Other tissues also showed several differentially expressed TEs, including thyroid, liver, skeletal muscle and adipose-subcutaneous (**Supplemental Table 6**).

To understand how sex-associated TE expression changes might affect biological function in breast mammary tissues, we performed gene ontology analysis of the protein-coding genes that were nearest the sex-associated TEs (**Methods**). These were significantly enriched for cellular components specific to stress fiber (P < 6.78 x 10^-14^, binomial test) and actin filament bundle (P < 1.54 x 10^-14^, binomial test). Myoepithelial cells of breast are known to contain large amounts of actin stress fibers and these fibers act as major mediators of cell contraction and help in milk production (Pellegrin and Mellor 2007), and organization of these actin fibers has also been shown to be associated with breast cancer (Tavares et al. 2017) suggesting TEs regulatory potential.

Notably, the genes that we found the most correlated with sex-associated TEs in breast mammary tissue have been implicated in breast cancer. For example, ANKRD30A and GABRP, known to specifically expressed in individuals with triple-negative breast cancer (Mathe et al. 2016; Sizemore et al. 2014) are positively correlated with two different MIR elements, both up-regulated in females (Fig 4 C, D). ELF5, a key regulator of mammary gland alveologenesis, the epithelial-mesenchymal transition in mammary gland development and breast cancer metastasis (Chakrabarti et al. 2012), is positively correlated with an L2 element that is upregulated in females (Fig 4 E). YBX1, a gene known to be involved in transfer RNA mediated breast cancer progression (Goodarzi et al. 2015) is negatively correlated with another L2 element (Fig 4F). These results suggest a potential association between TEs expression in female breast development and cancer.

### Ethnicity-associated TE transcription is most prevalent in skin

The role of TEs in pigmentation has been recently revealed in British peppered moths, where the insertion of a TE increases the transcription of the gene *cortex*, eventually giving rise to industrial melanism (Van’t Hof et al. 2016). Within humans, it has been shown that different ethnic groups express different genes associated with disease-phenotypes (Melé et al. 2015; Spielman et al. 2007) and that the expression of these genes is regulated by neighboring non-coding regions (Cheung et al. 2005). Most of the human studies, however, were either performed in cell lines or focused on gene expression, and little is known about TE expression across tissues in different human ethnic groups.

Here, we specifically investigated differential TE expression between 77 African American (AA) and 455 European American (EA) (**Supplemental Figure. 5)**. We used a similar linear regression approach as before. (**Methods**). We identified 614 subfamilies (468 increased in EA, 146 increased in AA) that showed significant differential expression (adjusted P-value < 0.05) between EA and AA (Fig. 5A, **Supplemental Table 7**).

**Figure 5.**
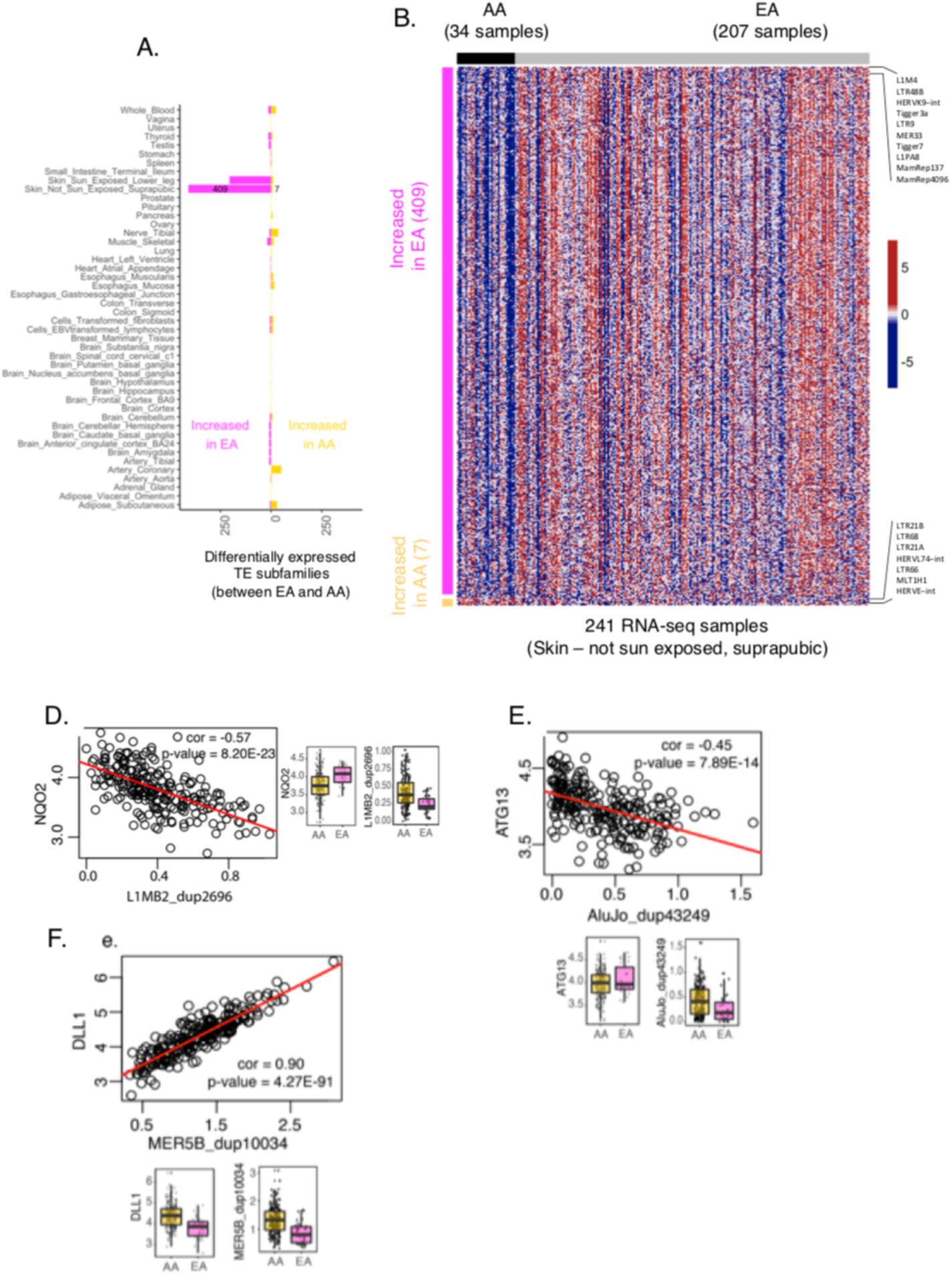
Differentially expressed TE subfamilies between ethnicities. (**A**) Double-sided stacked bar plot represents the number of differentially expressed TE subfamilies between EAs (European Americans) and AAs (African Americans) across different tissues TEs the expression of which was comparatively increased in EAs are shown on the left side, and those with expression increased in AAs on the right. The highest number of differentially expressed TE subfamilies were found in skin-not sun exposed tissue (416, log2 fold-change > 0 or < 0, adjusted P-value < 0.05) and most of them were increased in EAs. (**B**) Heatmap represents the expression of differentially expressed TE subfamilies in skin-not sun exposed tissue (red is higher and blue is lower). 416 subfamilies were shown in rows and 241 skin-not sun exposed samples in columns. Bar with the dark purple color on the left side of the heatmap represents a group of subfamilies that were up-regulated in EAs and the one with the gold color represents that were increased in AAs. The black/grey bar on the top of the heatmap identifies the AAs and EA samples. (**E-F**) Examples of ethnicity-associated TEs that were correlated with ribosome-biogenesis associated protein-coding genes in the skin (not sun-exposed). Scatterplots showing the TE expression (log2) on X-axis and gene expression (log2) on Y-axis, and the red line is the regression line. Each scatterplot is further separated into two (gene and TE) individual boxplots, where log-transformed expression was plotted in AAs and EAs. AAs were shown in gold and EAs were shown in purple colors. (**D**) For example, NQO2 was negatively correlated with a LIMB2_dup2696 copy in skin (not sun-exposed) tissue where NQO2 gene expression is higher in EAs and L1MB2_dup2696 is higher in AAs. (**E, F**) Similar plots were drawn using other top correlated TE and ribosome-biogenesis gene pairs.

At element level, we identified 17,119 differentially expressed TEs (**Supplemental Table 7**). Similar to sex-associated TEs, the effect size effect was small (log2 fold-change < 1) (Fig. 5A).

Skin (not sun-exposed) exhibited the largest number of differentially expressed TEs (416 subfamilies, 5,699 elements); the 409 subfamilies were increased in EAs, whereas brain regions showed the lowest number of differentially expressed TEs (Fig 5A, B, **Supplemental Table 7**). While there were also ethnic differences in TE expression in sun-exposed skin, the number of TEs differentially expressed here were about half of those in unexposed skin, suggesting that exposition to sunlight may somehow reduce differences in skin physiology between EA and AA.

Protein-coding genes near ethnicity-associated TEs in unexposed skin were significantly enriched in the various biological process, notably ribosome biogenesis (P < 1.89 x 10^-15^, binomial test). It has been shown that mutations in ribosomal proteins lead to dark skin (McGowan et al. 2008). Gene ontology analyses also revealed significantly enriched mouse phenotypes associated with the morphology of epidermis and keratinocytes, suggesting a possible role for TEs in skin morphology.

Notably, the topmost correlated genes were known to be involved in the maintenance of keratinocytes or in melanosome degradation. For example, NQO2, a gene known to limit skin carcinogenesis by protecting cells against oxidative stress (The Cancer Genome Atlas Network et al. 2012) is negatively correlated with an L1 element, which is up-regulated in AA (Fig. 5D) and ATG13, a known autophagic modulator in melanosome degradation (Murase et al. 2013), is also negatively correlated with an Alu element, upregulated in AA. It has been shown that EA skin-derived keratinocytes have more autophagic activity than those derived from AAs skin, suggesting a possible role for TEs in autophagic activity (Fig. 5E). Among the top positive correlated cases, we found a few genes involved in skin cell maintenance.

These include DLL1 known to regulate differentiation and adhesion cultured human keratinocytes (Estrach et al. 2008), which is positively correlated with a MER5B element upregulated in AA (Fig. 5F). These results suggest that TEs expression could underlay phenotypic differences between ethnicities, especially in the skin.

### Age-associated TE transcription is most prevalent in tibial artery

Unlike sex and ethnicity, several studies have investigated the association between aging and TEs in various species. For example, it has been shown that L1 elements were up-regulated in aging mouse liver (De Cecco et al. 2013b) and LINEs were up-regulated in the aging fly brain (Li et al. 2013). In addition, the fly study reported that the misregulation of TEs expression leads to age-dependent memory impairment and shortened lifespan. It also has been shown that Alu elements are up-regulated in patients with geographic atrophy (Kaneko et al. 2011) (age-related retinal pigment degeneration) and L1 elements in patients with Rett syndrome (Marchetto et al. 2010) (a rare genetic disease linked to developmental and nervous system problems). However, to our knowledge, the association of TE expression with age across many human adult tissues has not been investigated.

To detect age-associated changes in TE expression, we used a similar linear regression as before except age was treated as a continuous variable (**Methods**). We used individuals with 19 years to 70 years of age (**Supplemental Figure. 6 A, B)**. Overall, we identified 742 subfamilies that showed a significant change in expression with age (adjusted P-value < 0.05), the vast majority of the increasing expression (679 increased, 63 decreased, Fig. 6A, **Supplemental Table 8**). At the element level, we identified 27,189 TEs that change TE expression with age (**Supplemental Table 8**).

**Figure 6.**
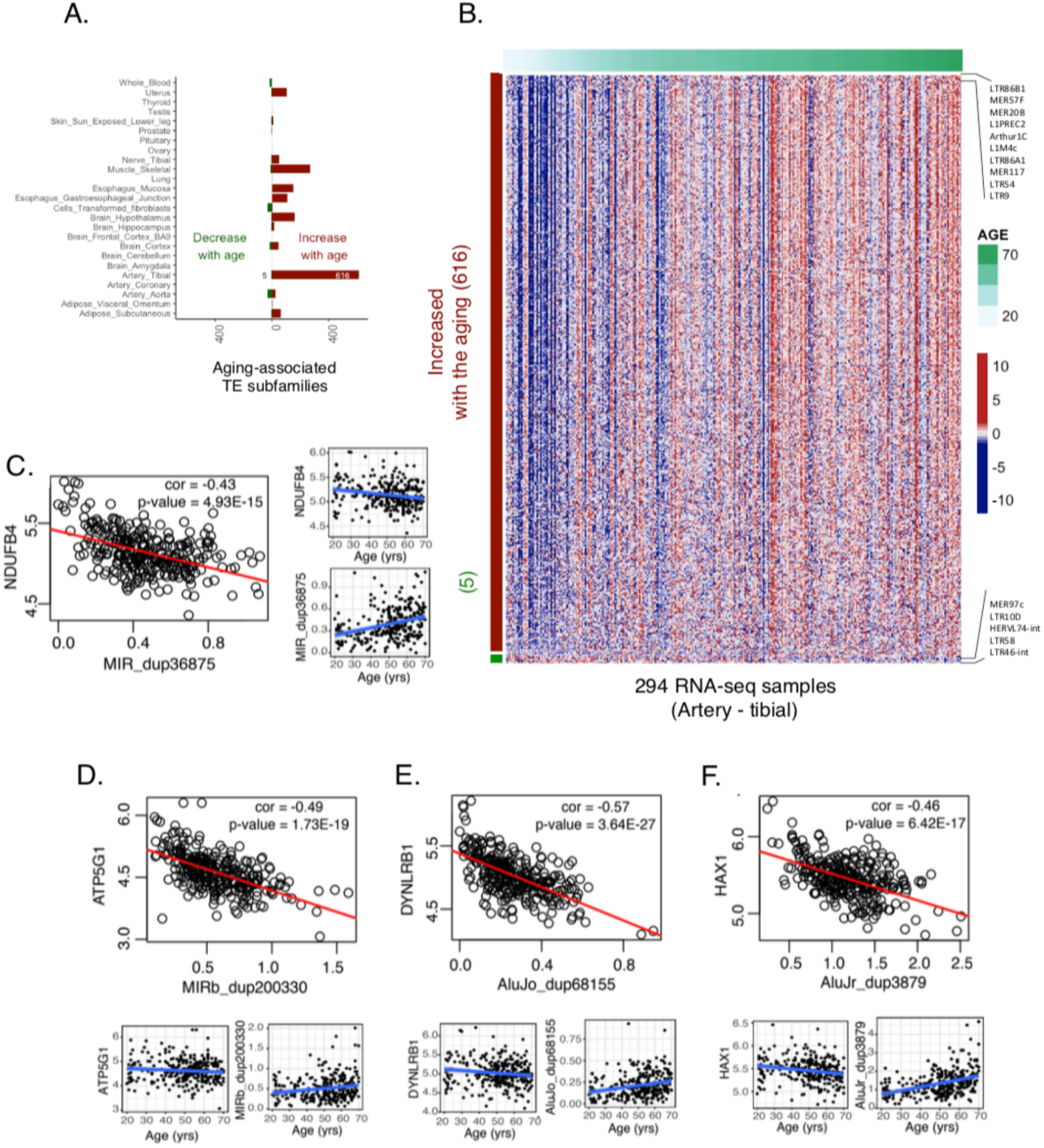
Aging-associated TE expression. (**A**) Double-sided stacked bar plot represents the number of age-associated TE subfamilies across different tissues. TEs the expression of which increased with the age are shown on the left side and those in which the expression decreased on the right. The highest number of TE subfamilies associated with aging were found in artery tibial tissue (631, log2 fold change > 0 or < 0, adjusted P-value < 0.05) and most of them were increased with the age. (**B**) Heatmap represents the expression of age-associated TE subfamilies in artery tibial tissue (red is higher and blue is lower). 631 subfamilies were shown in rows and 294 artery-tibial samples in columns. Artery-tibial samples were ordered from lower age-groups to higher. Red/green bar the left side of the heatmap represents identifies TE subfamilies increasing/decreasing expression with age. The green color bar on the top of the heatmap represents the age of the individuals. (**C-F**) Examples of Age-associated TEs that were negatively correlated with mitochondrial transport associated protein-coding genes in tibial artery tissue. Scatterplots showing the TE expression (log2) on X-axis and gene expression (log2) on Y-axis, and the red line is the regression line. Each scatterplot is further separated into two (gene and TE) individual scatterplots, where the log-transformed expression (on Y-axis) was plotted against age in years(on X-axis), and the blue line is the regression line. (**C**) For example, NDUFB4 was negatively correlated with a MIR_dup36875 copy in artery tibial tissue where NDUFB4 gene expression decrease with age and MIR_dup36875 increase with age. (**D-F)** Similar plots were drawn using another top negatively correlated TE and mitochondrial transport associated gene pairs.

There were a few tissues in which there was an association between expression of TEs and age, notably muscle, some brain regions, and a few others. However, the tissue that showest the largest number of age-associated TEs (621 subfamilies, 16,678 elements), nearly all increasing expression with age), was the tibial artery (Fig. 6B, **Supplemental Table 8**). The other artery-related tissues in GTEx including artery aorta and artery coronary did not show a similar trend. Atherosclerosis is a condition affecting arteries the incidence of which increases with age. A substantial number of histological images of the tibial artery in GTEx (145 out of 294) have been annotated by pathologists as being affected by this condition. Therefore, we have performed differential TE expression between affected and non-affected tibial artery samples, after controlling for age and other factors (**Methods**). We have detected two subfamilies (L1PA4 and LTR57) with increased expression in atherosclerotic arteries (**Supplemental Figure. 6C)**. Genes in the vicinity of some of the members of these families have been implicated in atherosclerosis: ACSL1 (Kanter et al. 2012), long-chain acyl-CoA synthetase 1, an enzyme known to be associated with accelerated atherosclerosis in diabetes and HPSE (Vlodavsky et al. 2013) is a mammalian enzyme that degrades heparan sulfate, a compound useful for arterial structures, and known to be associated with atherosclerosis. The other LTR57 member maps to a locus orthologous to a mouse locus in chromosome 8 associated to atherosclerosis (Burkhardt et al. 2011).

Protein-coding genes near age-associated TEs were significantly enriched for biological processes specific to cytokine production (P < 1.09 x 10^-59^, binomial test), response to type-1 interferon (P < 7.59 x 10^-31^, binomial test) and mitochondrial protein import activity (P < 4.7 x 10^-27^, binomial test). Notably, the topmost correlated genes were also known to be involved in either aging- or mitochondrial-related processes. For example, NDUFB4, a mitochondrial complex I subunit previously known to associate with the longevity of C.elegans (Yee et al. 2014) is negatively correlated with a MIR copy that increases in expression with age (Fig. 6C). ATP5G1, an inner mitochondrial membrane protein containing a mitochondrial targeting signal (Itakura et al. 2016) and also linked to aging (Houtkooper et al. 2011), is also negatively correlated with a MIRb copy that increases in expression with age (Fig. 6D). Genome-wide association studies in individuals with coronary artery disease found genetic risk loci in ATP5G1 and DYNLRB1 genes (Schunkert et al. 2011; Howson et al. 2017). DYNLRB1 is also negatively correlated with an Alu copy that increases in expression with age in the tibial artery (Fig. 6E). Among the top negatively correlated genes, many were associated with mitochondrial functions. For example, HAX1 is an anti-apoptotic protein that promotes mitochondrial fission and has been previously linked to the aging (Gatta et al. 2014); the expression of this gene is negatively correlated with an Alu copy (AluJr_dup3879), which increases in expression with age (Fig. 6F). Mitochondrial dysfunction and inflammation were some of the essential hallmarks of aging in mammals (López-Otín et al. 2013). Together, these results suggest that TEs might affect aging by regulating the protein-coding genes involved in mitochondrial or inflammation-related functions.

## Discussion

In this study, we have used the GTEx transcriptomic data across multiple tissue and individuals to provide a systematic and unbiased evaluation of transcriptionally active TEs in human adult tissues. Our results challenge the assumption that TEs are globally inactive in human adult tissues and support recent reports where a number of subfamilies were shown to be active (Faulkner et al. 2009; Fort et al. 2014). By greatly extending these reports, our study delineated a comprehensive atlas of transcriptionally active TEs across human adult tissues. We found that many TEs are tissue-elevated, often correlating, in a specific manner, with human phenotypes (breast regarding sex, skin regarding ethnicity, and tibial artery regarding age). Many TEs are positively correlated with nearby protein-coding genes. This could suggest a role of TEs in the regulation of gene expression, a role that is further supported by the finding that approximately 25% of non-coding expressed TEs contain eQTLs for nearby protein-coding genes.

Several recent studies have shown that using both uniquely mapped reads and multi-mapped reads to quantify TEs might introduce biases in expression (Lerat et al. 2017; Jin et al. 2015; Criscione et al. 2014). Another single-cell RNA-seq based study showed that integrating multi-mapped reads increases read coverage but not the overall results (Göke et al. 2015). To avoid this multi-mapping problem, we only used uniquely mapped reads to quantify TEs. Another technical issue is that often the entire TE containing regions are expressed and the TE expression in that region is not increased relative to the surrounding regions, usually hosting genes. Therefore, TE expression could just be a by-product off transcription of nearby genes. To address this issue, we built signal peaks from the RNA-seq reads and considered TEs expressed only if they correspond to one such RNA-seq peak (**Supplemental Figure 1A, Methods**). Thus, the TEs considered in our study is likely to be, at least partially, independent transcriptional events, and not the direct consequence of the transcription of nearby genes. Since these account for more than 100,000 of expressed TEs they dramatically affect downstream analyses.

Our study can be extended in several ways. First, GTEx is a collection of postmortem samples, and therefore transcription levels from these samples may be different from that of living individuals (Melé et al. 2015). Monitoring changes in TE transcription in living individuals would reveal unbiased results (Arda et al. 2016; Enge et al. 2017). Second, GTEx is not a disease associated project and most of the samples were collected from healthy individuals. However, TEs are known to be highly active in various types of cancer. Therefore systematically identifying TEs in different human cancer, or other disease-based, transcriptomic datasets could also reveal interesting insights. Finally, while using short sequence reads we have identified thousands of TEs, long-read sequencing technologies will enormously facilitate the mapping of repeat elements, leading to a much better characterization of the landscape of TE expression.

It has been shown that in mouse tissues several retrotransposable element subfamilies become transcriptionally active during normal aging, and further, that the expression culminates into active transposition in advanced age (De Cecco et al. 2013b). Our results show that expression of TEs increases with age in most tissues, supporting previous findings in the fly, mouse, and human (Li et al. 2013; De Cecco et al. 2013a, 2013b). Aging is a stochastic process where each cell in a tissue is affected in different ways, and analysis of adult tissues might be difficult to interpret (Enge et al. 2017). Most of the previous age-related transcriptome studies used different individuals with a different lifespan, and these studies could be prone to a bias that comes from individual genetic variation (Enge et al. 2017; Arda et al. 2016; Yang et al. 2015). Measuring samples from the same individual at different ages would reveal unbiased results that can help to understand potential markers of aging. Whether TE transcription is a cause or a consequence of aging-associated genomic alterations rather than the cause, requires additional data and experiments.

In summary, by resolving a longstanding problem whether TEs are globally silenced in humans or not, our study provides a comprehensive catalog of transcriptionally active TEs in human adult tissues, as well as their associations with a number of phenotypes. This catalog represents a valuable resource to the genomic, aging and other fields.

## Methods

### Data acquisition

Reads mapped to hg19 / GRCh37 for available mRNA sequencing data (n=8,555) together with matching phenotype covariates like sex, ethnicity, and age were downloaded on 20th December 2016 from the dbGAP data portal (data release V6, dbGaP accession phs000424.v6.p1). Briefly, reads were mapped by using TopHat (version 1.4.1) (Kim et al. 2013), parameters: --mate-inner-dist 300 --mate-std-dev 500 --no-sort-bam --no-convert-bam --GTF Homo_sapiens_assembly19.gtf --transcriptome-index. We used only the reads that mapped to unique locations in the genome (NH:i:1) to quantify TE expression. Individuals with genotyping issues, chromosomal or sex abnormalities were removed from the analysis, and this gave 8,051 RNA-seq samples in total. We downloaded repeats from RepeatMasker (UCSC Table Browser, hg19) and selected the ones that belong to transposable elements classes (LINE, SINE, LTR, and DNA), and Gencode genes (GV19, comprehensive annotation) from Gencode https://www.gencodegenes.org/.

### Expression quantification

We used featureCounts (version 1.4.3 from the subread package (Liao et al. 2014)) to count the number of uniquely mapped reads overlapping the transposable elements. Uniquely mapped reads were further normalized by the total number of reads in each tissue sample as RPKM. We considered a TE to be expressed if it satisfies the following steps.

1. TE with a normalized expression value greater than 0.1 RPKM in at least one sample. This step removes the noisy TEs with either lower number of uniquely mapped reads or expressed in a few samples.
2. TE that overlaps RNA-seq peak by at least one base pair in at least one sample (RNA-seq peak dataset was built by running MACS (Feng et al. 2012) default parameters) across every GTEx sample and pooling all the results or significant peaks into one) (**Supplemental Figure 1A**). This step removes noisy TEs that have an equal number of overlapping RNA-seq reads compared to their neighboring genomic regions.
3. TEs that overlap either protein-coding exons or proximal to the protein-coding promoter within 3 kb distance (Gencode v19 protein-coding gene annotation).

To estimate the fraction of the transcriptome mapped to the TEs, we divided the total number of reads mapped to each individual genomic TE element with the total number of mapped reads (x 10^6). To estimate the number of individual genomic TE copies expressed in each tissue, we counted the ones that have greater than or equal to 0.1 RPKM expression in at least one sample in each tissue separately.

### Tissue specificity estimation

TissueEnrich package (https://github.com/Tuteja-Lab/TissueEnrich) is used to calculate enrichment of tissue-elevated TEs. Tissue-elevated subfamilies are defined using the algorithm from the HPA (Uhlén et al. 2016), and grouped as follows:

1. Tissue Enriched: subfamilies with an expression level greater than 1 (normalized expression) that also have at least 1.5 fold higher expression levels in a particular tissue compared to all other tissues.
2. Group Enriched: Subfamilies with an expression level greater than 1 (normalized expression) that also have at least 1.5 fold higher expression levels in a group of 2-7 tissues compared to all other tissues, and that is not considered tissue-enriched.
3. Tissue Enhanced: Subfamilies with an expression level greater than 1 (normalized expression) that also have at least 1.5 fold higher expression levels in a particular tissue compared to the average levels in all other tissues, and that are not considered tissue-enriched or group-enriched.

Charlie6 subfamily was ignored from the analysis as it has less than one normalized expression in all samples.

### Estimation of correlation between TEs and nearest protein-coding genes in tissues and individuals using multivariate multilevel modeling

Gene and repeat expression define a bivariate vector. To decompose the raw variances and covariances of such vector into individual and tissue components a multivariate multilevel model was adopted (Snijders 2011). Assuming a nesting structure; measurement nested within individuals, and individuals within tissues. This leads to three levels: measurements within individuals within tissues. The first level is that of the dependent variables indexed by *h* = *1,2*, the second level is that of the individuals *i = 1,…,n_j_*, where *n_j_* denotes the number of individuals in *j*-th tissue, and the third level is that of the tissues, *j = 1,…,N*, where *N* denotes the number of tissues. The multivariate model is formulated as a hierarchical linear model using dummy variables used to indicate the dependent variables

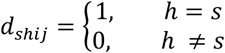

With these dummies, the random intercept models for the dependent variables can be integrated into one three-level hierarchical linear model (Raudenbush, S.W. and A.S. Bryk. 2002) by the expression

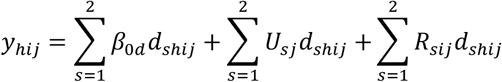

In particular, the notation denotes: *y_1ij_*gene expression from individual *i* at tissue *j*, *y_2ij_*repeat expression from individual *i* at tissue *j*, *β_01_* intercept of gene expression, *β_02_* intercept of repeat expression, *U_1j_* the random effect at tissue level of gene expression, *U_2j_* the random effect at tissue level of repeat expression, *R_1ij_* the random effect at the individual level of gene expression *R_2ij_* the random effect at the individual level of repeat expression. We assume:

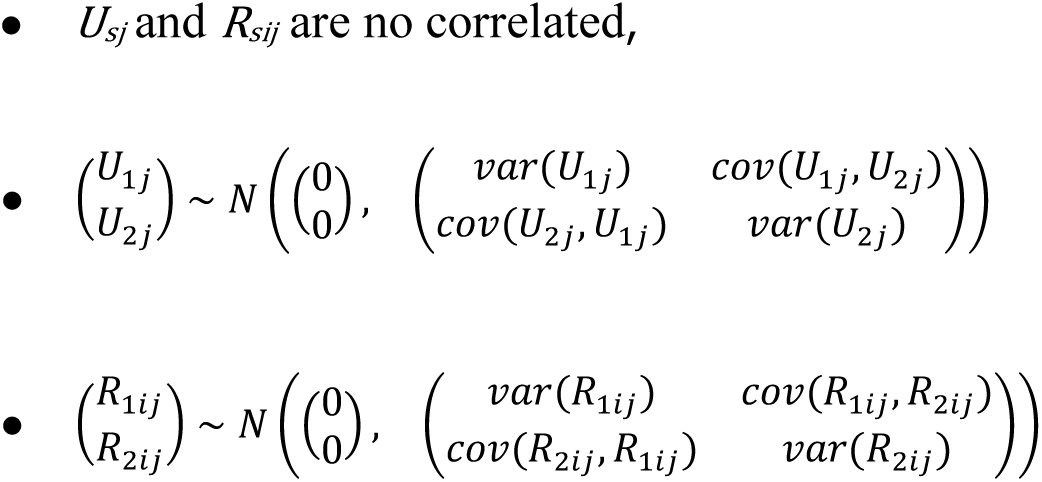

Model parameters were estimated using lme function implemented in the nlme package from R. Then, correlation coefficients at the tissue and individual level were computed from variances and covariances estimates. Notice that GTEx data shows cross-classification structure. Individuals are not perfectly nested in tissues. Weighting methods to address this fact will be implemented in further investigations.

### eQTL analysis

We downloaded eQTLs from dbGAP data portal (data release V6, dbGaP accession phs000424.v6.p1). We only considered significant cis-eQTLs (1 Mb distance to the transcription start site) with a nominal p-value is less than nominal p-value threshold (nominal p-value threshold for calling a variant-gene pair significant for the gene). In total, there were around 26 million SNP-gene pairs (unique SNPs = 2,552,394 and unique genes = 28,059). Next, we overlapped 79,558 non-coding TEs and found 19,944 (25%) overlapping at least one SNP. To calculate correlations, we used MVML method. We found 3,284 TEs correlated with nearest genes (out of 19,944 expressed TEs that overlap eQTLs); 3284 (3007 positive, 277 negative), and 36,532 TEs were correlated with nearest genes (out of 59,614 expressed TEs that do not overlap eQTLs); 36,532 (31,911 positive, 4,621 negative).

### Variance decomposition analysis

To assess the contribution of tissue and individual to TE expression variation, we used a linear mixed model (LMM). TE expression was modeled as a function of tissue and individual (considered as random factors). The LMM was implemented in the R package lme4 (Bates et al. 2015). To obtain the variance components, we divided the restricted maximum likelihood (REML) estimators for the random effects of tissue, individual and residual variance by their sum. TEs that were not expressed (RPKM > 0) in any one of the samples were excluded from the analysis. We used log2 (RPKMs) to normalize the data and pseudocounts to deal with zero expression values.

### Linear regression analysis

In each tissue, we modeled TEs expression using the following linear regression model (Ritchie et al. 2015):

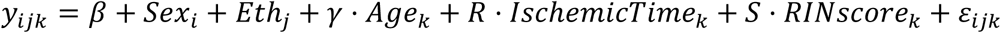 with *i* = 1, 2; *j* = 1, 2; *k* = 1, …, *n_ij_*, *n_ij_* is the number of samples in the (*i,j*)-th condition. *y_ijk_* denotes the expression level of a transposable element in the *i*-th level of sex, *j*-th level of ethnicity and *k*-th sample in the (*i,j*)-th condition. *β* denotes the regression intercept, *Sex_i_* denotes the effect of *i*-th level of Sex, *Eth_j_* denotes the effect of the *j*-th level of Ethnicity, *γ* denotes the regression coefficient of *Age*, *Age_k_* denotes the age of the *k*-th sample in the (*i,j*)-th condition, *R* denotes the regression coefficient of Ischemic time, *IschemicTime_k_* denotes the ischemic time of the *k*-th sample in the (*i,j*)-th condition and *S* denotes the regression coefficient of RIN score, *RINscore_k_* denotes the RIN score of the *k*-th sample in the (*i,j*)-th condition, and *ε_ijk_* denotes the error term of the (*i,j,k*)-th observation, that we assume is normally distributed. In addition, we selected the TEs with higher than 0.1 RPKM expression in at least 10 samples and log-transformed the data before applying linear regression.

### Sex-association analysis

We examined the expression of TEs in the non-coding regions of the human genome across 42 tissue-sites using an expression threshold > 0.1 expression in more than ten samples in each tissue that was shared between both sexes. This threshold enables more robust estimates of sex-associated TEs in the examined samples, but it excludes TEs with lower individual-level expression (<=10 individuals) and also excludes sex-specific tissues from the analysis. Samples used in this analysis were originated from 540 individuals of whom 186 were female, and 354 were male. The demographic details related to these samples are provided in (**Supplemental Fig. 3A**). While we found no significant differences in age at death (Mann-Whitney P-value = 0.32) between female and male samples, we did detect significant differences in the ischemic time (Mann-Whitney P-value < 0.05) and RNA integration score (RIN score) (Mann-Whitney P-value < 0.05) (**Supplemental Fig. 3B**). However, we verified that none of the findings reported in this study could be affected by these factors.

### Ethnicity-association analysis

We utilized GTEx data that belong to 77 EA and 455 AA ethnic groups. To keep at least a few samples in both groups across each tissue, we discarded Asian, American Indian or Alaska native or individuals with unknown groups (**Supplemental Fig. 4A, C**). We detected significant differences in the sex, age, ischemic time and RNA integration score (Mann-Whitney P-value < 0.05) between EA and AA ethnic groups (**Supplemental Fig. 4B**). However, we verified that none of the findings reported in this study could be affected by these factors.

### Aging-association analysis

We grouped individuals into five age-groups ranging from 19 to 70 years (**Supplemental Fig. 3A, B**). We used tissues with at least one samples across all these five groups. Like before, we also corrected aging regression for sex, ethnicity, ischemic time and RIN-score. If γ was significantly deviated from 0, TE was considered to be age-associated. TEs was up-regulated with age if *γ*> 0 and down-regulated if *γ* < 0. For atherosclerosis analysis, we annotated tibial artery samples using histo-pathological images. We found 145 samples out of 294 tibial artery samples have atherosclerosis annotation (in V6). We applied linear regression to calculate differential TE expression between tibial artery samples with and without atherosclerosis. All this analysis was done by correcting Gender, RIN score, Ischemic time and Age.

### Gene-ontology analysis

We used the GREAT tool (McLean et al. 2010) using the nearest gene approach with the whole genome as a background. For larger datasets, we selected “Significant By Region-based Binomial testing”.

## Data access

All the raw data can be accessed from https://www.ncbi.nlm.nih.gov/projects/gap/cgi-bin/study.cgi?study_id=phs000424.v6.p1

## Acknowledgements

M.A.M-R. acknowledges the support of the Spanish Ministry of Economy and Competitiveness (BFU2017-85926-P), Centro de Excelencia Severo Ochoa 2013-2017 (SEV-2012-0208) and AGAUR (SGR468).

## Author contributions

G.K.B performed all the analysis. F.R performed multivariate correlation analysis. G.K.B, R.G, and M.P.S designed the analyses, interpreted and wrote the manuscript. G.K.B and R.G conceived the project.

## Supplemental Figures

**Supplemental Figure 1. Filtering reliable TEs using RNA-seq peaks. (A)** TEs that overlap RNA-seq peak by at least one base pair in at least one sample was used in this study (RNA-seq peak dataset was built by running MACS default parameters across every GTEx sample and pooling all the peaks into one). This step removes noisy TEs that have an equal number of overlapping RNA-seq reads compared to their neighboring genomic regions. (**B**) For example, THE1D-int_dup861 (THE1D-int subfamily / ERVL-MaLR family / LTR class) on chr6 (start-129024007, end-129025585) expressed in heart tissues was on the many identified by using the RNA-seq peak filter (highlighted in yellow color). (**C**) another example (chr21:33054843-33054978: MIRb subfamily / MIR family / SINE class) with no RNA-seq peak (highlighted in blue)

**Supplemental Figure 2. Correlation between TEs and nearest protein-coding genes. (A)** Distribution of correlation between TEs and their nearest protein-coding genes in tissues and individuals. (**B**) Significant correlation differences (Wilcoxon test, *** (P < 0.001)) between TEs that overlap eQTLs and nearest protein-coding genes, and the ones that do not overlap eQTLs and nearest genes. Median correlation scores were shown in each group (all – black, positive pairs – in blue, negative pairs – in red).

**Supplemental Figure 3. Group and tissue-enriched TE subfamilies. (A, B)** The network of the group- and tissue-enriched subfamilies across different tissues. Central nodes are tissues and outer nodes are repeats. Central nodes in orange represent group-enriched and in blue represent tissue-enriched. Directed edges connect repeats (outer nodes) with tissues (central nodes).(**C**) Boxplots represent the group or tissue-enriched expression of various subfamilies across different tissues. log2 normalized expression was shown on the y-axis and tissue names of the x-axis. LTR75B is a group-enriched subfamily specific to different brain tissue types, and LTR88c and LTR28 are tissue-enriched subfamilies specific to pancreas and pituitary tissues respectively. (**D**) Gene ontology analysis of TEs with highest tissue variance

**Supplemental Figure 4. Distribution of different sex-groups of GTEx individuals. (A)** Summary of the demographic details of the individuals in this study separated by sex. Values of age (in years), ischemic time (in hours) and RIN (RNA integration) score displayed as a median across male and female individuals and “No” represents the number of individuals. (**B**) Boxplots showing the distribution of the age, RIN score and ischemic time male and female individuals. Differences between male and female samples were shown as N.S (Not Significant) or significant (Mann-Whitney U test, p-values, ** = 0.01, *** = 0.001).

**Supplemental Figure 5. Distribution of different ethnic-groups of GTEx individuals. (A)** Summary of the demographic details of the individuals in this study separated by two different ethnic groups, EA (European American) and AA (African American). Values of age (in years), ischemic time (in hours) and RIN (RNA integration) score displayed as a median across EA and AA ethnic groups and “No” represents the number of individuals. (**B**) Significant differences between age, ischemic time and RIN score between EAs and AAs (Mann-Whitney U test, p-values, ** = 0.01, *** = 0.001). (**C**) Distribution of individuals separated by the ethnic group across different tissues.

**Supplemental Figure 6. Distribution of different age-groups of GTEx individuals. (A)** Number of individuals across five different age groups from 19 years to 70 years. (**B**) Distribution of the number of individuals separated by tissue and age group.

## Supplemental Tables

**Supplemental Table 1.** Transcriptionally active TEs in human tissues. It can be downloaded from https://public-docs.crg.es/rguigo/Data/gbogu/Repeat_paper_data/111222_tes_8051_samples.matrix.main.ann.backup.rds

**Supplemental Table 2.** List of TEs overlapping eQTLs.

**Supplemental Table 3.** Global correlation between TEs and nearest genes in tissue and individuals.

**Supplemental Table 4.** Variation of TE expression in tissues and individuals.

**Supplemental Table 5.** Tissue-elevated transposable elements in human adult tissues.

**Supplemental Table 6.** Differentially expressed TEs between males and females.

**Supplemental Table 7.** Differentially expressed TEs between European American and African American individuals.

**Supplemental Table 8.** Differentially expressed TEs between individuals of different age groups.

